# Firing behavior of single motor neurons of the tibialis anterior in human walking as non-invasively revealed by HDsEMG decomposition

**DOI:** 10.1101/2022.04.03.486869

**Authors:** Hikaru Yokoyama, Naotsugu Kaneko, Atsushi Sasaki, Akira Saito, Kimitaka Nakazawa

## Abstract

Investigation of the firing behavior of spinal motor neurons (MNs) provides essential neuromuscular control information because MNs form the “final common pathway” in motor control. The MNs activated during human infants’ leg movements and rodent locomotion, mainly controlled by the spinal central pattern generator (CPG), show highly synchronous firing. In addition to spinal CPGs, the cerebral cortex is involved in neuromuscular control during walking in human adults. Thus, MN firing behavior during adult walking is expected to be similar to that of infants and rodents and has some unique features. Recent technical advances allow non-invasive investigation of MN firing by high-density surface electromyogram (HDsEMG) decomposition. Therefore, we investigated the MN firing behavior of the tibialis anterior muscle during walking by HDsEMG decomposition. We found motor unit recruitment modulation compared with steady isometric contractions, doublet firings, and gait phase-specific firings during walking. We also found high MN synchronization during walking over a wide range of frequencies, probably including cortical and spinal CPG-related components. The amount of MN synchronization was modulated between the gait phases and motor tasks. These results suggest that the central nervous system, including the spinal CPG and cerebral cortex, flexibly controls MN firing to generate appropriate muscle force during human walking. In addition to revealing the neural control mechanisms of walking, our data demonstrate the feasibility of non-invasive investigation of MNs during walking, which will open new frontiers for the study of neuromuscular function in medical and exercise sciences.

## Introduction

Each spinal motor neuron (MN) innervates multiple muscle fibers, forming a motor unit (MU), the smallest organizational component of the neuromuscular system. Thus, MU activity provides fundamental information on neuromuscular control (Farina et al., 2014). The central nervous system regulates the muscle contraction force required for various movements by changing MN activity, such as the firing rate, recruitment, and degree of synchronization of firings (Heckman and Enoka, 2012).

Spinal central pattern generators (CPGs), which produce rhythmic locomotor activity, have essential roles in generating MN activity during walking (Kiehn, 2016). In neonatal rodents, MNs in a muscle fire synchronously, on the order of milliseconds, via electrical gap junctions in CPGs (Chang et al., 1999). Likewise, gap junctions in adult mammalian CPGs contribute to the production of locomotor behavior (Kiehn and Tresch, 2002). Such highly synchronized MN activation may be crucial for efficiently generating coordinated motor patterns (Kiehn and Tresch, 2002). Although human infants have limited motor abilities, CPGs allow them to generate primitive locomotor-like movements (Dominici et al., 2011). A recent study revealed highly synchronized MN firing during leg movements in human infants (Del Vecchio et al., 2020). Although locomotor (-like) movements in human infants and rodents are mainly controlled by the CPGs, both the CPGs and the cerebral cortex significantly contribute to the neuromuscular control in human adult locomotion (Petersen et al., 2012; Yokoyama et al., 2019). Therefore, MN firing behavior during human adult walking may partially follow the knowledge previously reported in human infants and rodents but has some unique features.

Although MU firing has been traditionally investigated in humans using the decomposition of intramuscular electromyographic (EMG) signals based on a template-matching algorithm (De Luca et al., 1996), intramuscular EMG has considerable limitations due to its invasiveness. A previous study attempted to investigate MN firing by applying a template-matching algorithm to surface EMG (sEMG) signals measured during walking (De Luca et al., 2015). Because this study focused on the feasibility of the approach, the details of MN firing behavior during walking remain largely unclear. Additionally, doubts have been raised concerning the validity of sEMG decomposition based on template matching (Farina et al., 2015; Enoka, 2019).

Recently, another EMG decomposition approach that uses high-density sEMG (HDsEMG) and blind source separation (BSS) algorithms has been widely used (Farina et al., 2017; Hug et al., 2021; Liu et al., 2021). Originally developed for isometric muscle contractions, this method has also been utilized on non-isometric muscle contractions such as hand movements (Chen et al, 2020), ankle dorsiflexion (Oliveira and Negro 2021), and infant leg reflexes (Del Vecchio et al, 2020). Regarding the feasibility of using HDsEMG decomposition on dynamic muscle contractions, a recent simulation study demonstrated that HDsEMG decomposition is applicable even during non-isometric contractions of the tibialis anterior (TA) muscle, up to an ankle dorsiflexion range-of-motion (ROM) of 30°, less than that of slow walking (Yokoyama et al., 2021).

Therefore, we here investigated MN activity of the TA during walking using HDsEMG decomposition. We hypothesized that: (1) significant MN synchronization would be observed in the beta band (15–40 Hz) in addition to the frequency band lower than 5 Hz; (2) doublets (a pair of spikes at short intervals less than 15 ms) would be observed; and (3) MN recruitment patterns would differ between the stance and swing phases and between walking and ankle isometric contraction. The first hypothesis is based on the fact that spinal CPGs cause low-frequency band MN synchronization (Del Vecchio et al., 2020) and cortical descending drives are involved in beta band MN synchronization (Farina et al., 2014). The second hypothesis is derived from evidence that doublets were observed during mammalian locomotion (Gorassini et al., 2000) and the high commonality of locomotor neural networks among species (Grillner, 2011). The third hypothesis is based on task specificity of MU recruitment (Hudson et al., 2019) and functional differences in TA activity during the swing and stance phases (Perry, 1992).

## Materials and Methods

### Participants

Thirteen healthy men (aged 22–42 years) with no history of neural or musculoskeletal disorders participated. Each provided written informed consent for participation in the study, which was performed in accordance with the Declaration of Helsinki and approved by the Ethics Review Committee of The University of Tokyo (approval number: 701-2).

### Experimental setup and design

The participants walked on a treadmill (Bertec, Columbus, OH, USA) at 0.6 m/s for 10 minutes. The last nine minutes of the data were collected. A relatively slow walking speed was chosen considering the effects of ankle joint ROM and movement artifacts on HDsEMG decomposition (Mentiplay et al., 2018; Yokoyama et al., 2021). A previous simulation study demonstrated that HDsEMG decomposition analysis is applicable even during non-isometric contraction of the TA muscle up to an ankle dorsiflexion ROM of 30°, and the decomposition accuracy is equivalent to that during an isometric contraction when ankle ROM is lower than 20° (Yokoyama et al., 2021). During slow walking, ankle ROM is about 20° up to 0.6 m/s, increases sharply above 0.6 m/s, and reaches about 30° at comfortable walking speed (1.0–1.2 m/s) (Mentiplay et al., 2018). In addition to the joint angle changes, the signal quality of the EMGs also affects the HDsEMG decomposition. The number of motor units identified by HDsEMG decomposition is highly sensitive to the signal quality (Holobar et al., 2009). Noise contamination caused by the movement of the recording electrodes, cabling, and lower limb soft tissues during locomotion must be minimized, and the severity depends on walking speed (Guidetti et al., 1996).

The participants also performed an isometric dorsiflexion task in the sitting position. The ankle angle was fixed at 0° using an ankle-foot orthosis. Participants performed the maximal voluntary contraction (MVC) task for approximately 3 s twice. The highest value was assumed to be the MVC. After the MVC task, they performed two submaximal isometric contractions at a 25% MVC, which was sustained for 30 s and had a rising phase of 5 s (Fig. 1). The root mean square (RMS) value of the EMG signals was displayed as visual feedback of the contraction level, and the participants were asked to keep following the target contraction level, which was also displayed concurrently with the participant’s contraction level. All tasks were performed with sufficient rest for at least three minutes.

**Figure 1.**
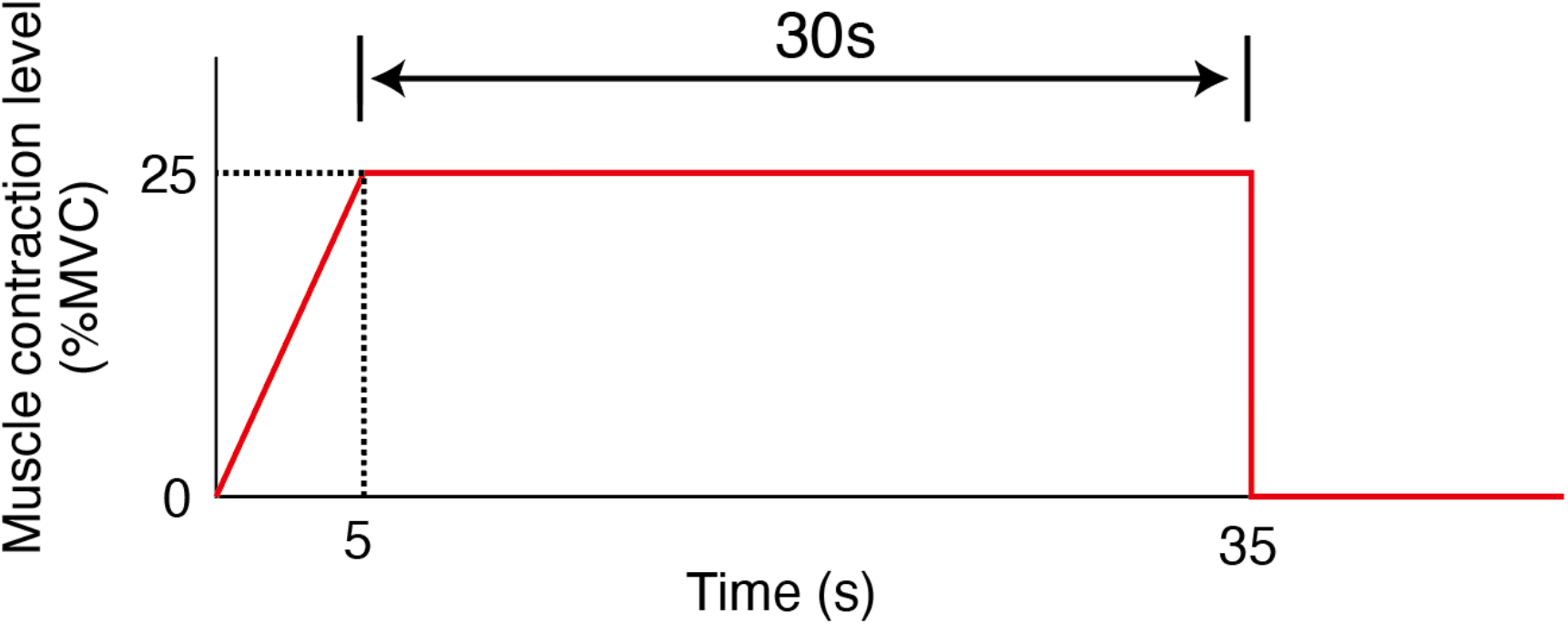
Time course of the muscle contraction level in the isometric contraction task. Red line indicates the muscle contraction level that participants are required to follow. MVC: maximum voluntary contraction.

### Data collection

HDsEMG signals were recorded from the right TA using a two-dimensional adhesive grid of 13 × 5 equally spaced electrodes (each 1 mm in diameter, inter-electrode distance 8 mm), with one electrode absent from the upper right corner (GR08MM1305, OT Bioelettronica, Torino, Italy). The hair was shaved the day before or the day of the experiment, and the skin was wiped with sanitary cotton moistened with alcohol. An experienced operator determined the perimeter of the superficial region of the muscle by palpation and marked its profile on the skin. The grid electrode was attached to the marked area, arranged longitudinally along the muscle and fixed to the skin using adhesive tape. A reference electrode was placed over medial surface of the right tibia. This position was selected because it is less likely to be affected by joint movements. The EMG recording setup is illustrated in Fig. 2. All the equipment for EMG recording (e.g., electrodes, cables, wireless amplifier, and wireless trigger signal receiver) was fixed with an elastic underwrap on the tibia to minimize movement artifacts. Electrode-skin impedance was reduced using conductive paste (Elefix, NIHON KOHDEN, Tokyo, Japan). HDsEMG signals were recorded in monopolar form and converted to digital data using a 16-bit wireless amplifier (Sessantaquattro, OTBioelettronica, Torino, Italy). The signals were amplified 256 times, sampled at 2000 Hz, and band-pass filtered between 10 and 500 Hz. The EMG signal was recorded directly onto a MicroSD card to prevent data loss, using the data logger mode of the Sessantaquattro system. Before HDsEMG decomposition analysis, 64 monopolar EMG signals were re-referenced to form 59 bipolar signals by taking the difference between adjacent electrodes in the direction of the longitudinal axis of the muscle.

**Figure 2.**
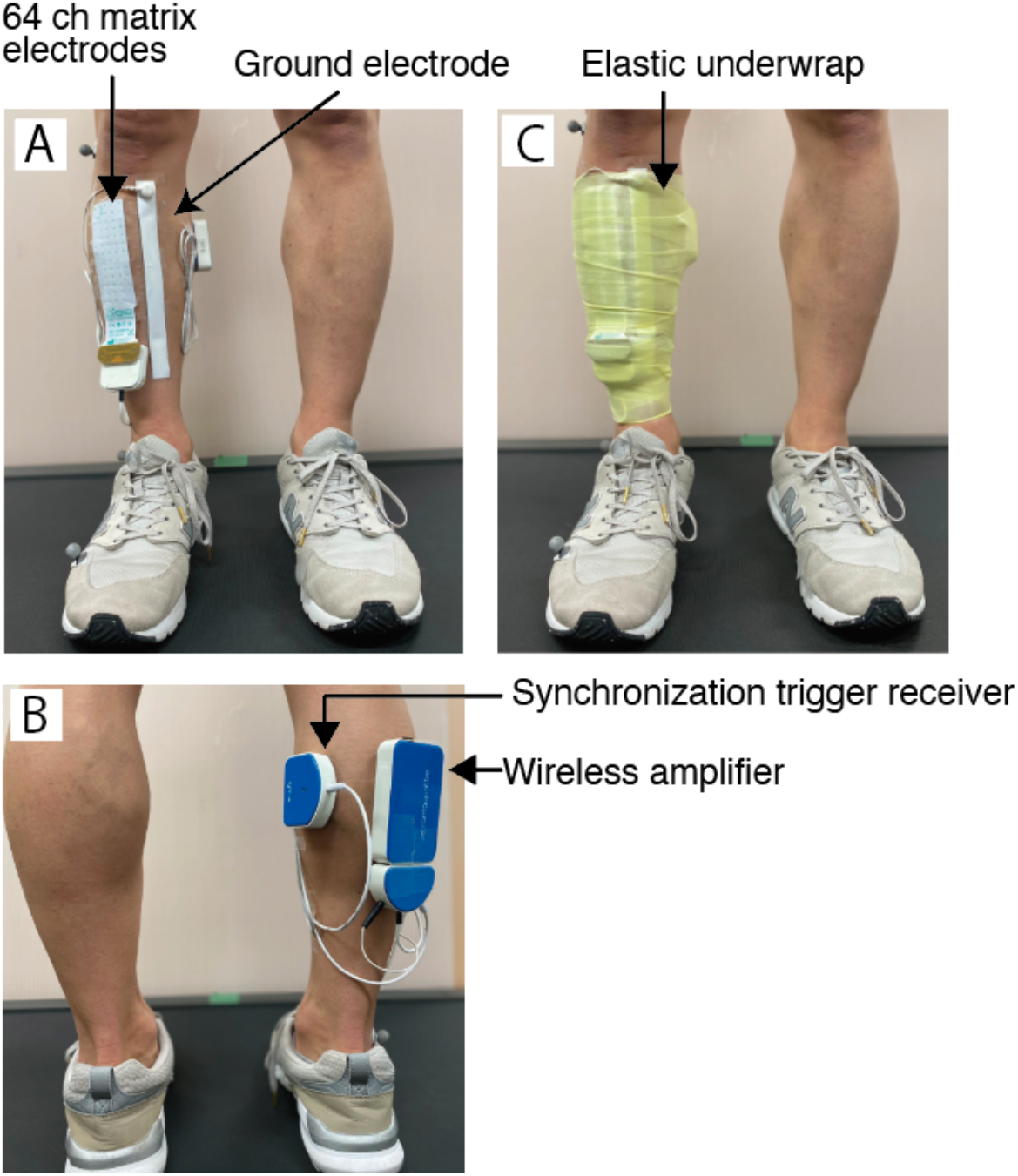
The setup of wireless high-density surface EMG (HDsEMG) measurement. (A) Front view of the setup before fixing with elastic underwrap. (B) Back view of the setup before fixing with elastic underwrap. (C) Front view of the setup after fixing with elastic underwrap.

Three-dimensional ground reaction forces were recorded from force plates under the right belt of the treadmill at 1000 Hz during the walking task.

Kinematic data were recorded at 100 Hz during the walking task using an optical motion capture system composed of 11 cameras (OptiTrack system with Flex3 cameras, Natural Point, OR, USA). Three spherical markers were placed on the right side of the fifth metatarsal head (toe), lateral malleolus (ankle), and lateral femoral epicondyle (knee).

### Data analysis

#### Kinematic and kinetic data analysis

The kinematic signals (marker location data) were digitally smoothed with a zero-lag low-pass Butterworth filter (6-Hz cutoff, fourth-order). The ankle joint angle was calculated from the marker coordinates. The ground reaction force data were smoothed using a low-pass filter (5-Hz cutoff, zero-lag Butterworth filter, fourth-order). The heel-contact (HC) timing was detected based on the vertical component of the ground reaction forces (threshold: 20 N).

#### HDsEMG decomposition

HDsEMG signals were decomposed into single motor unit activity using an automatic convolutive BSS technique based on fastICA and the convolutional kernel compensation approach (Negro et al., 2016). HDsEMG decomposition was performed independently for HDsEMG signals in the walking and submaximal isometric contraction tasks. The quality of decomposition was evaluated using a silhouette (SIL) value (Negro et al., 2016). The SIL is the difference between the within- and between-cluster sums of point-to-centroid distances, normalized by dividing by the maximum of the two values. In this study, MUs with SIL >0.87 were used for further analyses (Hassan et al., 2019; Negro et al., 2020).

#### Characteristics of motor units

We examined the firing rate of MNs based on inter-spike intervals (ISI). A pair of motor unit action potentials with short inter-spike intervals (typically 5–15 ms) are called “doublets” (Thomas et al., 1999; Mrówczyński et al., 2015). In rodents, the motor units of the TA exhibit doublets during walking (Gorassini et al., 2000), indicating their significant role in enhancing the muscle output force. Therefore, doublets are considered a functional strategy of the central nervous system to improve muscle contraction efficiency during motor tasks that require a large force. Doublets were defined as two consecutive discharges with an ISI of 15 ms or less (Thomas et al., 1999). We regarded MNs that showed doublet discharges when doublets were detected in >30% of the steps. We evaluated the frequency of occurrence of doublets in non-overlapping 15 ms bins from 300 ms before to 300 ms after heel contact and from 300 ms before to 300 ms after toe-off.

We compared the MU recruitment patterns between walking and isometric contraction tasks. We first examined the recruitment threshold of each MU based on the amplitude of the rectified and low-pass filtered EMG signals (Butterworth, fourth-order). Considering the muscle contraction speed, the cut-off frequency was set at 1 Hz for the isometric contraction task and 4 Hz for the walking task (Yavuz et al., 2018; Yokoyama et al., 2019). The recruitment threshold was obtained separately in each trial for the two isometric contractions, and in every stride for walking. We then identified the same MUs used in both the walking and isometric contraction tasks using the MU tracking method (Martinez-Valdes et al., 2017, 2018; Oliveira and Negro, 2021). For this analysis, the representation of the motor unit action potentials (MUAPs) in the two dimensions of the identified MUs was extracted by spike-triggered averaging based on the spike times and crosscorrelations among MUs (Martinez-Valdes et al., 2017, 2018; Oliveira and Negro, 2021). We regarded only MUs with high peak cross-correlation (r > 0.80) as the same MUs across tasks (Cogliati et al., 2020; Oliveira and Negro, 2021). Using the tracked MUs, we calculated the correlation coefficient of the recruitment threshold between isometric contraction and walking. We also performed a correlation analysis between the first and second trials to verify the reliability of the recruitment thresholds obtained from the small number of trials during isometric contraction.

During walking, the TA muscle shows two activity peaks: (1) in the early swing phase (phase 1) and (2) in the initial stance phase (phase 2). We examined whether MNs were recruited during a specific phase. The peak activity was higher in phase 2 than in phase 1. To remove the effects of the size principle, we examined the number of MUs that were activated only during phase 1. We defined phase 1-specific MNs as those whose discharge was detected in more than 30% of the steps during phase 1, but not during phase 2. We defined the period for the two phases as follows: (1) 100 ms before to 300 ms after the toe-off for phase 1 and (2) 100 ms before to 300 ms after the heel contact for phase 2.

#### Synchronization of activity of motor neurons

Because the neural connections of each MN in the pool vary even in an individual muscle, the synchronization strength of each MN does not represent the synchronization characteristics of the MN population caused by common inputs. Therefore, instead of evaluating synchronization between pairs of MNs, previous researchers estimated MN synchronization with pooled spike trains from multiple MNs in two independent groups with random permutation of MN members in each group (Del Vecchio et al., 2020; Liu et al., 2021). In this study, we randomly selected six MNs from those identified, divided them into two groups of three each, and summed the spike trains in each group. Because previous studies have used six MNs to reliably estimate coherence (Dai et al., 2017; Liu et al., 2021), the MN synchronization analysis was performed only in participants whose number of identified MNs was six or more in the two activation phases of TA during walking and the isometric contraction task. Eight of the thirteen participants were included in the analysis. Before the cross-correlation analysis, smoothing was applied with a convolution with a 10-ms Hann window to the motor unit spike trains (Del Vecchio et al., 2020). The window length was set to reflect the short-term synchronization between MNs within 10 ms (Reilly et al., 2004) in the crosscorrelation analysis with permutated and pooled MU spike trains. Cross-correlation was calculated independently in the two TA activation phases and isometric contraction task. In the walking task, spike trains were segmented based on the two activation phases of the TA. For each activation phase, we extracted data in the period, which was the same as that used for the detection of phase specificity, in each gait cycle and concatenated all extracted data across gait cycles. To use the same data length among the tasks and participants, we used 100-gait cycle data (40 s [100 × 400 ms segments]) for phase 1 and phase 2 during walking. For the isometric contraction task, we used 40 s of data composed of two intermediate 20 s data segments after removing the ramp-up period and the initial and last 5 s from the two trials. This procedure was iterated 100 times or number of times corresponding to possible combinations for dividing MNs into two groups (when the identified number of MNs was lower than 12) with permutations of the assigned MNs to the two groups.

We also evaluated MN synchronization in the frequency domain (i.e., coherent oscillations) among the MN spike trains. To examine frequency-domain synchronization, we used coherence analysis within the groups of motor units (as described above for the cross-correlation analysis). Coherence was calculated as follows:

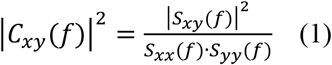

where *Sxy(f)* is the cross-spectrum between *x* and *y*, and *Sxx(f)* and *Syy(f)* are the auto spectra for *x* and *y*, respectively, for frequency *f*. The coherence was computed in 400 ms nonoverlapping windows and assigned MNs were permutated to two groups, as described for the time-domain cross-correlation analysis. We used unfiltered spike trains, the data before smoothing by convolution, in the cross-correlation analysis based on previous studies (Del Vecchio et al., 2020; Liu et al., 2021). The significance level of coherence is defined by the following equation:

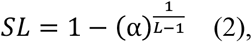

where α is the confidence level (0.05), and *L* is the number of segments (steps) used for the analysis (Rosenberg et al., 1989; Halliday et al., 1995)

#### Statistical analysis

The normality of the datasets was tested using the Kolmogorov-Smirnov test. The number of motor neurons was compared between walking and isometric contraction tasks using paired t-test. The number of motor neurons was also compared between phases 1 and 2 during walking using paired t-test.

Differences between the correlation coefficients regarding the MU recruitment threshold were tested using Fisher’s Z-transformation.

Peak values of the cross-correlation were compared between phases 1 and 2 during walking and the isometric contraction task using the non-parametric comparison method with 1000 random permutations (Nichols and Holmes, 2002) because normality was not observed in all cases. The p-value from the permutation test was corrected using false discovery rate (FDR) for multiple comparisons (Benjamini and Hochberg, 1995).

Coherence values were compared between phases 1 and 2 during walking and the isometric contraction task using the permutation test with FDR correction because normality was not observed in all cases. Statistical tests for coherence values were separately performed in the following four frequency bands: delta (2.5–5 Hz), alpha (7.5–10 Hz), low-beta (12.5–20 Hz), and high-beta (22.5–40 Hz). Although the alpha band is usually defined as between 8–12 Hz, in this study, we defined its frequency between 7.5–10 Hz because of the low-frequency resolution of the coherence analysis (2.5 Hz resolution) and an observed frequency peak at 12.5 Hz in many participants, which, when extended to 15 Hz, seemed to reflect a low-beta band component.

The firing rate of MN discharges was also compared between phases 1 and 2 during walking and the isometric contraction task using the permutation test with FDR correction because normality was not observed in all cases.

## Results

### Ankle-dorsiflexion range of motion

Mean ankle-dorsiflexion ROM across participants was 20.8° (standard deviation [SD]: 2.3°, range: 15.7–24.1°). The ankle-dorsiflexion ROM in this study was within the applicable range for the HDsEMG decomposition suggested by a simulation study (Yokoyama et al., 2021).

### Motor unit activity identified from HDsEMGs

Figure 3 shows examples of EMG signals recorded from the TA during walking. Noise levels of the recorded EMG signals were low based on visual inspection.

**Figure 3.**
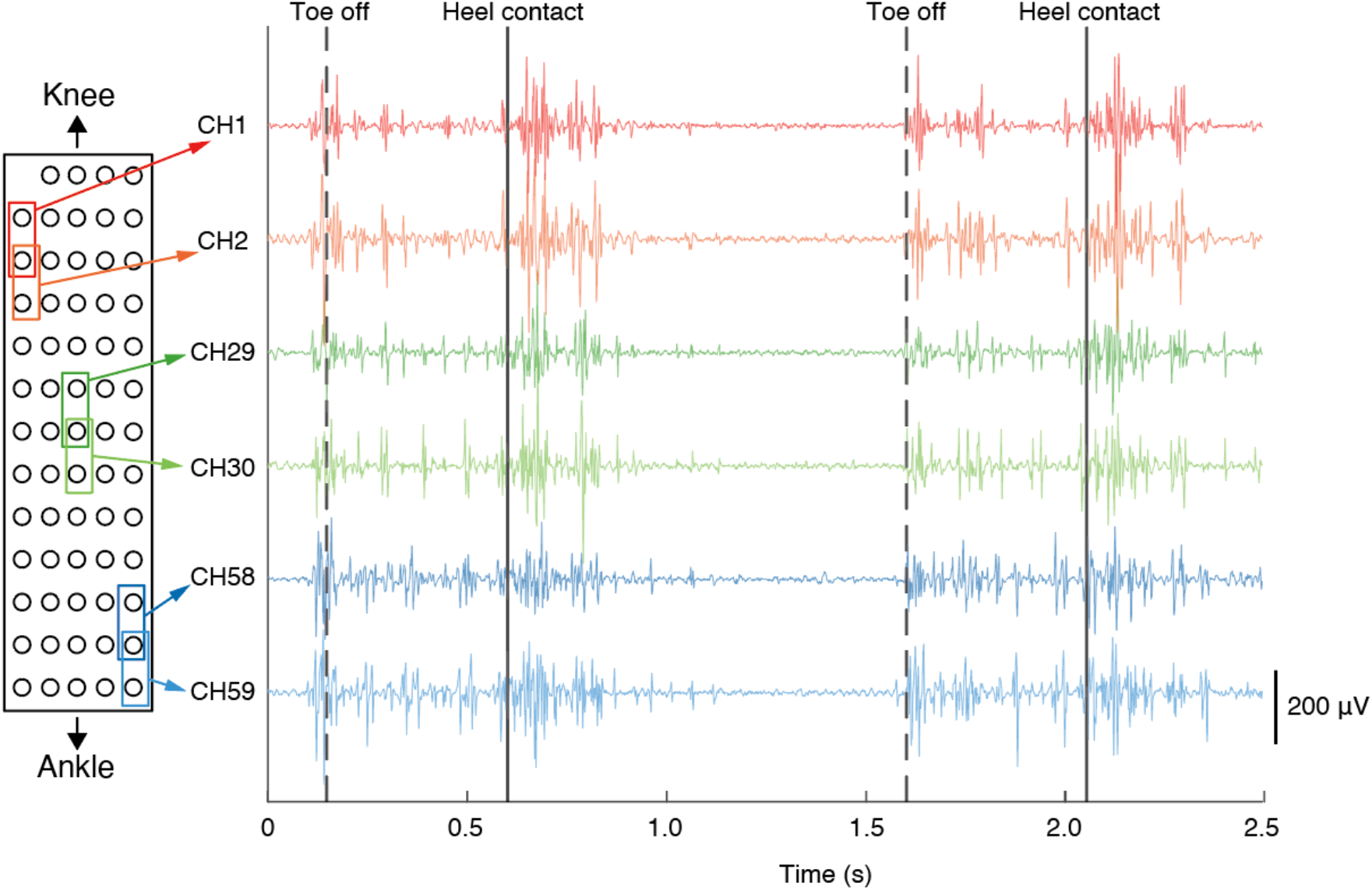
Examples of recorded EMG signals of the tibialis anterior muscle during walking. The left side shows the electrode arrangement used in this study. The surrounding rectangles highlight pairs of electrodes for each represented differential-signal channel. Solid and dashed lines indicate heel contact and toe-off timings, respectively.

Figure 4A shows the grand-averaged activation pattern of the TA in a gait cycle across participants. TA activity showed two activation phases during walking. The first activation phase was from immediately before the toe-off to the mid-swing phase (phase 1). The second phase was from just before heel contact to mid-stance (phase 2). Figure 4B shows an example of a time series of muscle activation and MN discharge of the TA from a participant. MN discharges were observed during the two abovementioned phases of TA activity. MN3 in this example exhibits doublets, as highlighted by the dark blue rectangles in Fig. 4B. Some MNs fired only during phase 1 (MN5) or phase 2 (MN8 and MN9).

**Figure 4.**
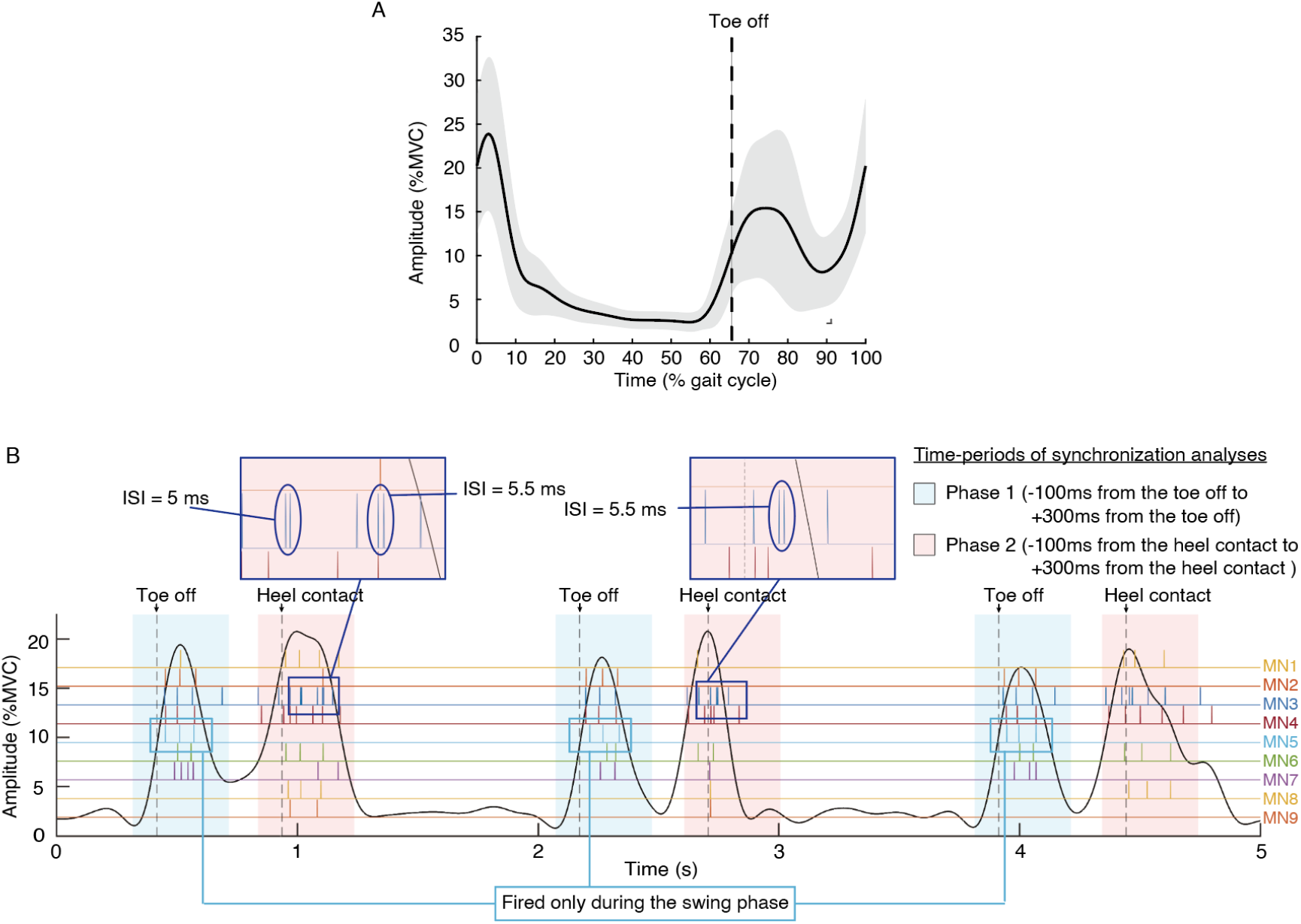
Muscle activation pattern and spike trains of motor neurons of the tibialis anterior (TA) muscle during walking. (A) Averaged activation pattern of the TA in a gait cycle across participants. The gray area indicates the standard deviation. The vertical dashed line represents the timings of toe-off. (B) Example of a participant’s muscle activation and motor neuron discharges of the TA. Typical examples of the doublets (two consecutive discharges with very short interspike intervals (ISI)) are zoomed up for better observing in the deep blue rectangles. Spike trains are represented as binary signals (high value: firing timing). The vertical dashed lines represent the timings of heel contact or toe-off. The blue- and red-shaded areas show the time periods used for coherence analyses of the TA muscle’s first and second activation peak, respectively.

The number of identified MNs was significantly higher in the isometric contraction task (21.3±7.3 [mean±SD]) than in walking (13.4±5.2) (Fig. 5A, p = 3.5×10^-6^, paired t-test). We also compared the number of identified MNs between the two phases of TA activity during walking. The number of identified MNs was significantly higher in phase 2 (12.3±5.7) than in phase 1 (7.5±4.6) (Fig 5B, p = 0.0084, paired t-test).

**Figure 5.**
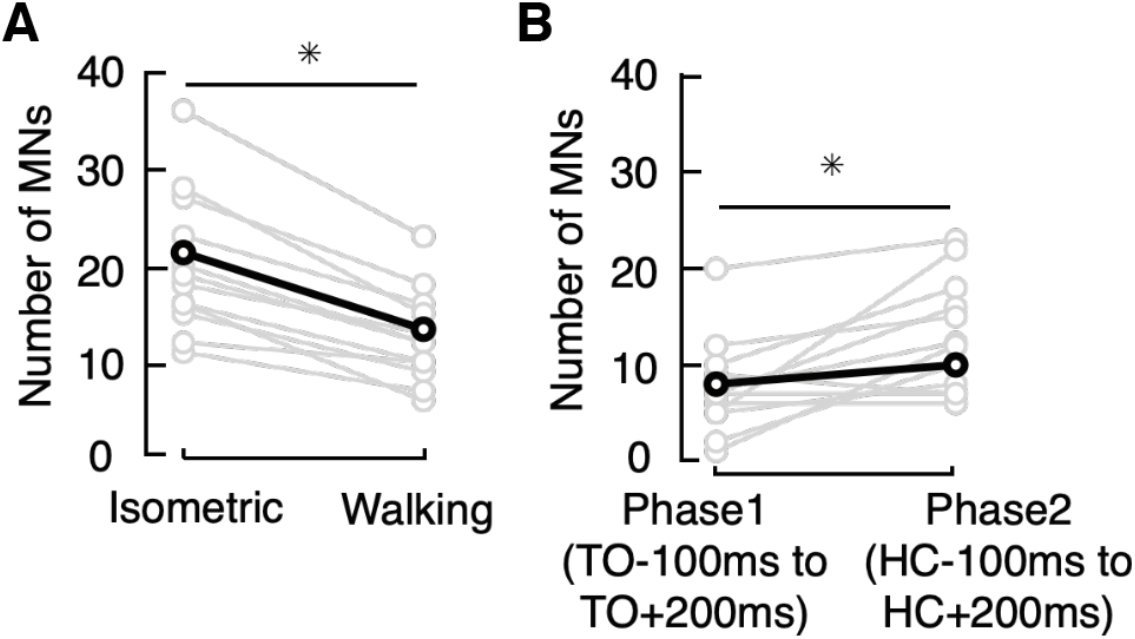
Identified number of motor neurons (MNs). (A) Comparison of the number of MNs between the isometric ankle dorsiflexion task and walking. *:p<0.05 (paired t-test). (B) Comparison of the number of MNs between two walking phases corresponding to the two peak timings of the TA activation during walking. TO: toe-off, HC: heel contact. *:p<0.05 (paired t-test).

Table 1 summarizes the activation phase specificity of MN recruitment and doublets of MN discharges. Regarding phase specificity, because the muscle activity level was lower in phase 1 than in phase 2 (Fig. 4A), we only focused on phase 1-specific MNs considering the effects of the size principle. Phase 1-specific MNs were observed in 5 out of 13 participants (38.5%). Across MNs identified in all participants, 16 out of 174 MNs (9.20%) were phase 1-specific.

**Table 1.** Gait phase specificity and doublets of motor neuron (MN) activities.

|  | Rate within participants | Rate of appearance<br>within all MNs across participants |
| --- | --- | --- |
| Phase 1-specific MNs | 5/13 (38.5%) | 16/174 (9.20%) |
| MNs which exhibited doublets | 10/13 (76.9%) | 33/174 (19.00%) |

MNs that exhibited doublets were observed in 10 out of 13 participants (76.9%) (Table 1). Across MNs identified in all participants, 33 of 174 MNs (19.0%) exhibited doublets (Table 1). During phase 1 of TA activity, doublets were frequently observed at the following two timings: (1) around the toe-off (about 50 ms before to 25 ms after toe-off) and (2) the initial swing to mid-swing (about 125 ms to 225 ms after toe-off) (Fig. 6A and 6C). During phase 2, doublets were frequently observed during the initial contact with the loading response (0 to 100 ms after heel contact) (Fig. 6B and 6D).

**Figure 6.**
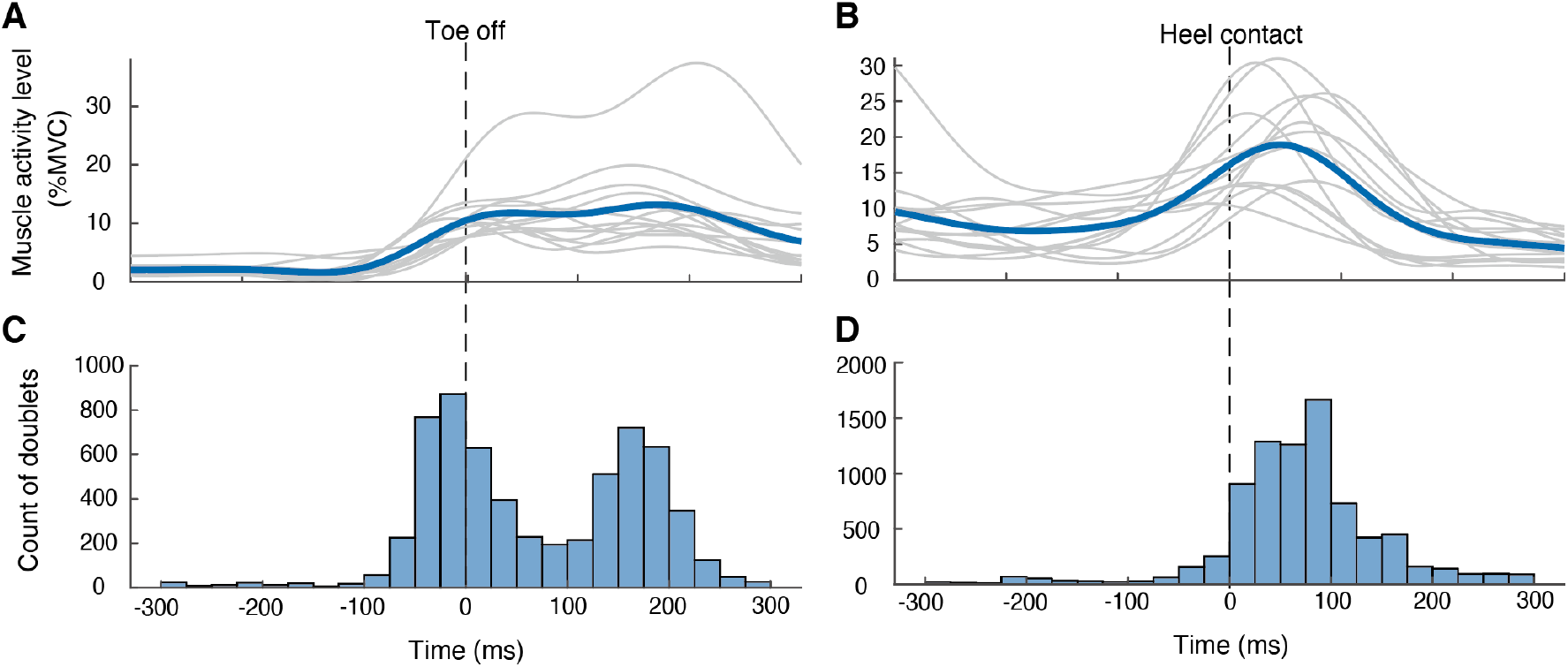
Muscle activation patterns and frequency of occurrence of doublets in the tibialis anterior muscle during walking. (A, B) Muscle activation patterns of TA around toe-off and heel contact. Blue lines indicate the average patterns across participants. Gray lines represent individual data. (C, D) Histograms of timing of doublets around toe-off and heel contact. Vertical dashed lines represent toe-off or heel contact timings.

One hundred eight MUs were tracked between the walking and isometric contraction tasks out of 174 MUs identified in the walking tasks. An example of the similarity between the MUAPs of a tracked MU is shown in Figure 7A. The MU recruitment thresholds between the two trials of the isometric contraction task were highly correlated (r = 0.82; Fig 7B). In contrast, the correlation of recruitment thresholds between the first trial of the isometric contraction task and the walking task was weak (r = 0.27, Fig 7C). The two correlation values were significantly different (p < 0.001, Fisher’s Z test).

**Figure 7.**
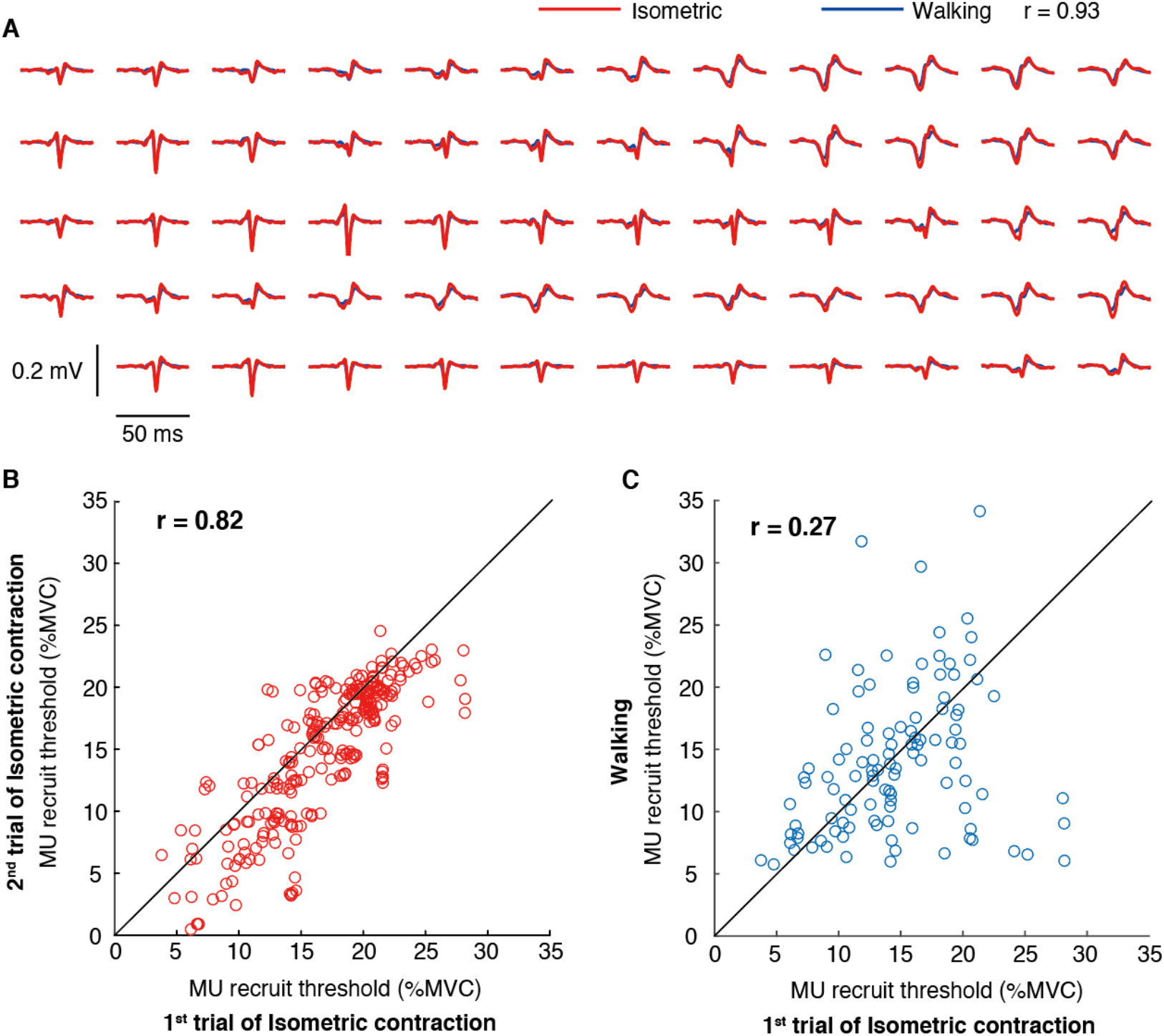
Matching motor units from isometric contractions and walking and motor unit recruitment thresholds. (A) An example of matched motor unit action potential (MUAP) shapes of each electrode between the isometric contraction and walking tasks. (B, C) Relationship of the motor unit (MU) recruitment threshold. The relationship between the first and second trials of the isometric contraction tasks is shown in panel B, and that between the first trial of the isometric contraction and walking tasks is shown in panel C. Each plot indicates individual MU data. The identity lines are provided in the two plots for reference.

### Motor neuron synchronization

Figure 8A shows the mean and individual data of the time-domain cross-correlation analysis in three conditions (the two phases of TA activity during walking and the isometric contraction task). The peak value of the cross-correlation was consistently nearly 0 among the participants (Fig. 8A). The mean peak values were significantly different among all conditions (p = 0.0078 for all comparisons between the two conditions, permutation test with FDR correction, Fig. 8B).

**Figure 8.**
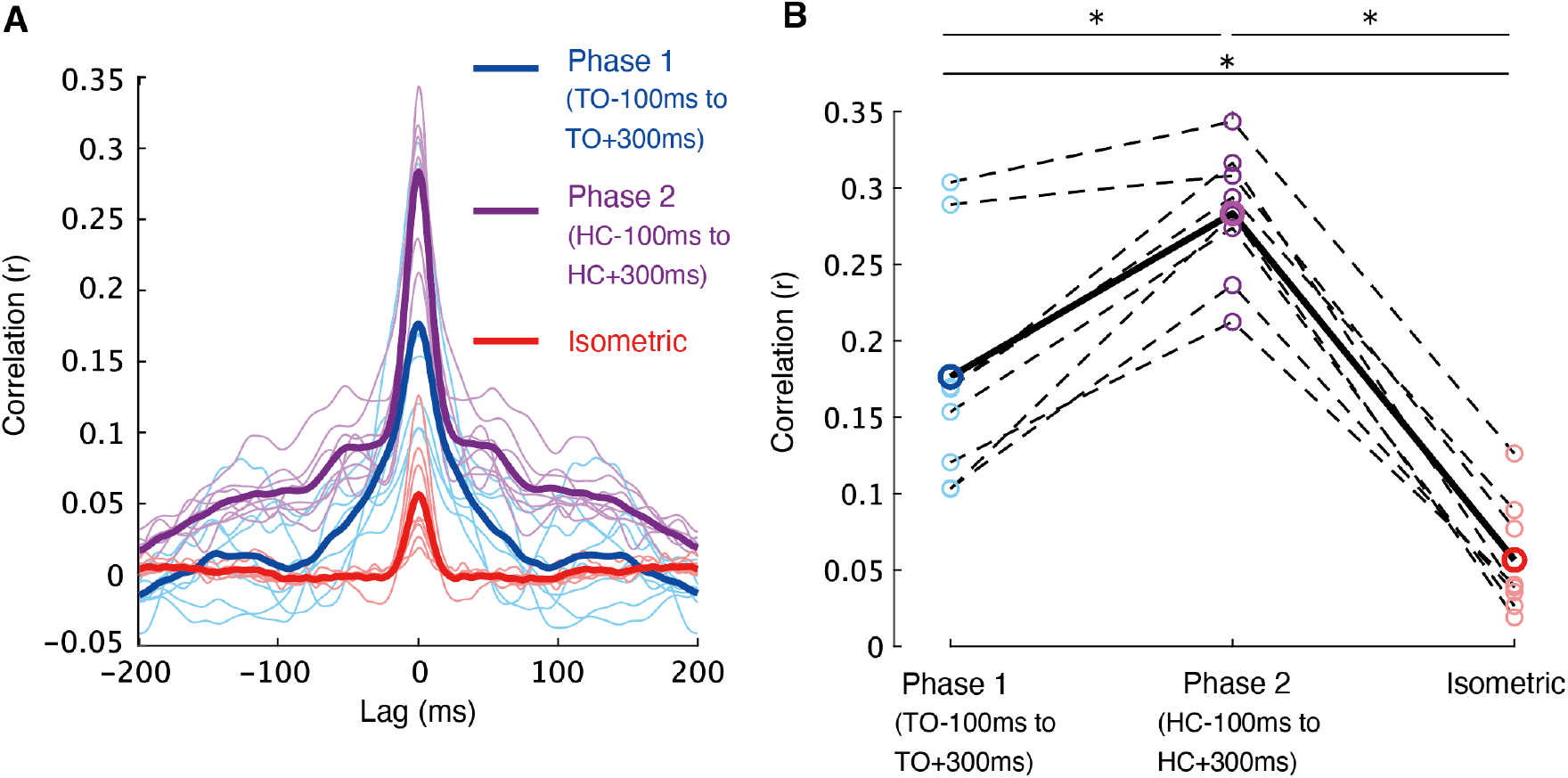
Cross-correlation of motoneuron activities. (A) Mean (thick lines) and individual (thin lines) cross-correlation between two sets of motor neurons’ activities in the two walking phases and the isometric ankle dorsiflexion task. TO: toe-off, HC: heel contact. (B) Peak of the cross-correlation. Mean (solid line with thick-lined circle) and individual (dashed line with thin-lined circle) data are shown. *:p<0.05 (permutation test with FDR correction).

We further examined the synchronization inputs to the MN pools in the frequency domain by using coherence analysis (Fig. 9). Representative data from three participants (Figs. 9A1–9A3) and averaged data across participants (Fig. 9D4) for the three conditions are shown. We found significant coherence in the wider frequencies below 50 Hz in all conditions, especially below 5 Hz (significance threshold = 0.0298, see Methods). In phase 1 during walking, high coherence between 10–20 Hz peaking at 12.5–15 Hz was a unique feature compared to the other two conditions. In phase 2 and the isometric contraction conditions, instead of the 10–20 Hz coherent oscillation component, a clear 20–40 Hz (extended to 50 Hz in some participants) coherent oscillation component was observed. The coherent oscillation component higher than 20 Hz was particularly evident in phase 2 during walking. Next, we compared the strength of coherence among the conditions in four frequency bands: delta (2.5–5 Hz), alpha (7.5–10 Hz), low-beta (12.5–20 Hz), and high-beta (22.5–40 Hz) (Fig. 8B). In the delta band, the coherence value was significantly higher in the order of phase 2 of walking, phase 1 of walking, and isometric contraction (p = 0.031 [phase 1 vs. phase 2], 0.011 [phase 1 vs. isomeric contraction], and 0.011 [phase 2 vs. isometric contraction], permutation test with FDR correction). In the alpha band, the coherence value was significantly higher in phase 2 than in isometric contraction (p = 0.29 [phase 1 vs. phase 2], 0.082 [phase 1 vs. isomeric contraction], and 0.023 [phase 2 vs. isometric contraction]; permutation test with FDR correction). In the low-beta band, the coherence values in the two walking phases were significantly higher than those in isometric contraction (p = 0.94 [phase 1 vs. phase 2], 0.023 [phase 1 vs. isometric contraction], and 0.023 [phase 2 vs. isometric contraction], permutation test with FDR correction). In the high-beta band, the coherence value was significantly higher in phase 2 than in phase 1 and isometric contraction (p = 0.012 [phase 1 vs. phase 2], 0.18 [phase 1 vs. isomeric contraction], and 0.012 [phase 2 vs. isometric contraction]; permutation test with FDR correction).

**Figure 9.**
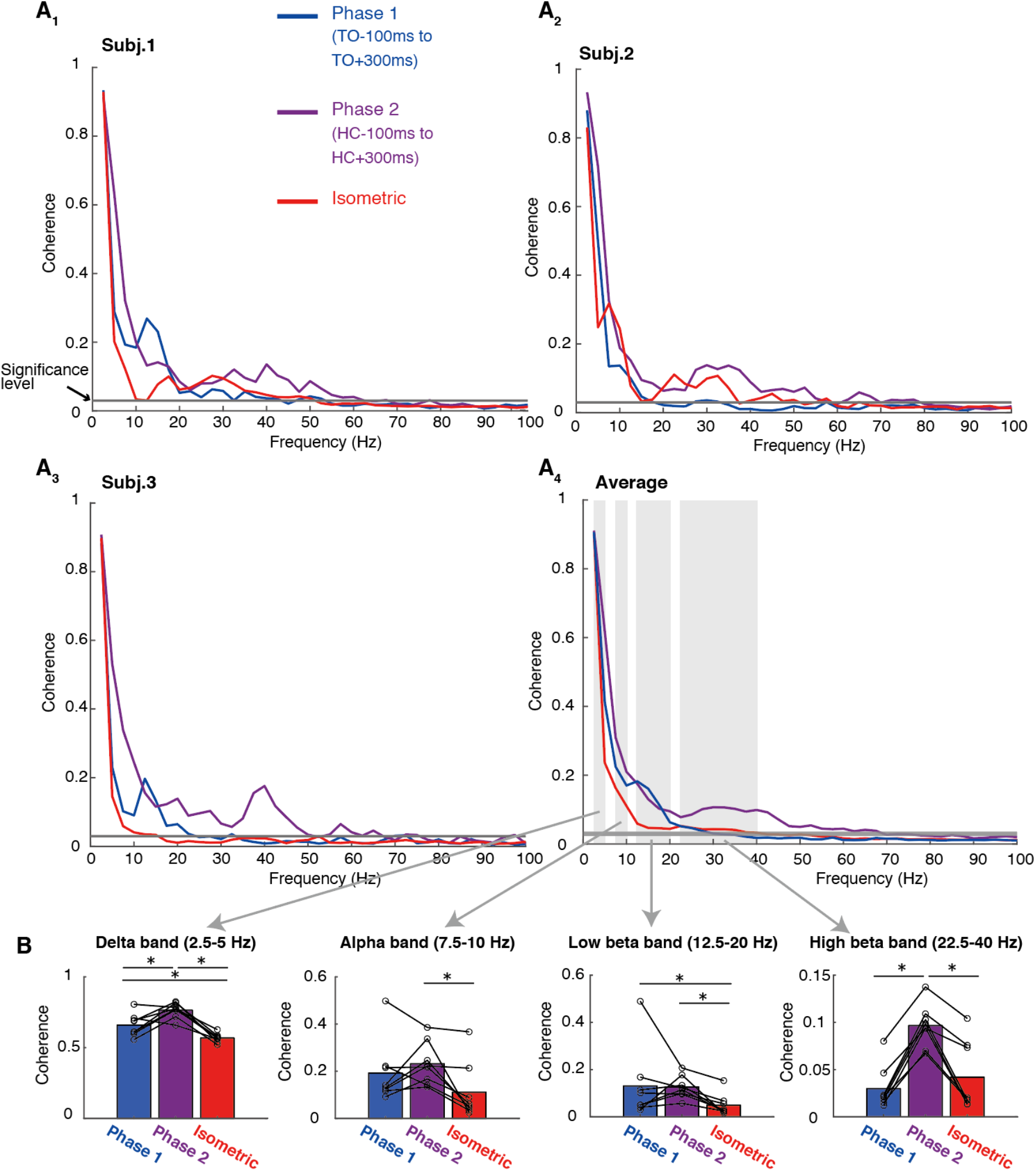
Coherence of motor neuron activities. (A) Coherence between two sets of motor neurons’ discharges in the two walking phases and the isometric contraction task. Example data from three participants (A1-A3) and averaged data across participants (A4) are also shown. Gray horizontal lines indicate a significant threshold for coherence value. Gray shaded areas correspond to frequency bands used for comparisons among the conditions. (B) Comparisons of coherence values among the two walking phases and the isometric task in four frequency bands. Bar plots represent mean values across participants and line plots represent individual values. *:p<0.05 (permutation test with FDR correction).

Previous studies have shown that the estimated degree of synchronization between spiking neurons is positively correlated with the firing rate of MN populations (Vecchio et al., 2019). Therefore, we compared the firing rates of the MNs in the three conditions from the participants used for the synchronization analysis (Fig. 10). The MN firing rates during the isometric condition (mean ± SD: 14.1 ± 2.5 spikes/s) was not significantly different from that of phase 1 (13.0 ± 1.41 spikes/s) and phase 2 (15.9 ± 3.3 spikes/s) during walking (p = 0.26, isometric vs. phase 1; p = 0.07 for isometric vs. phase 2, permutation test with FDR correction), but that between phases 1 and 2 was significantly different (p = 0.047, permutation test with FDR correction).

**Figure 10.**
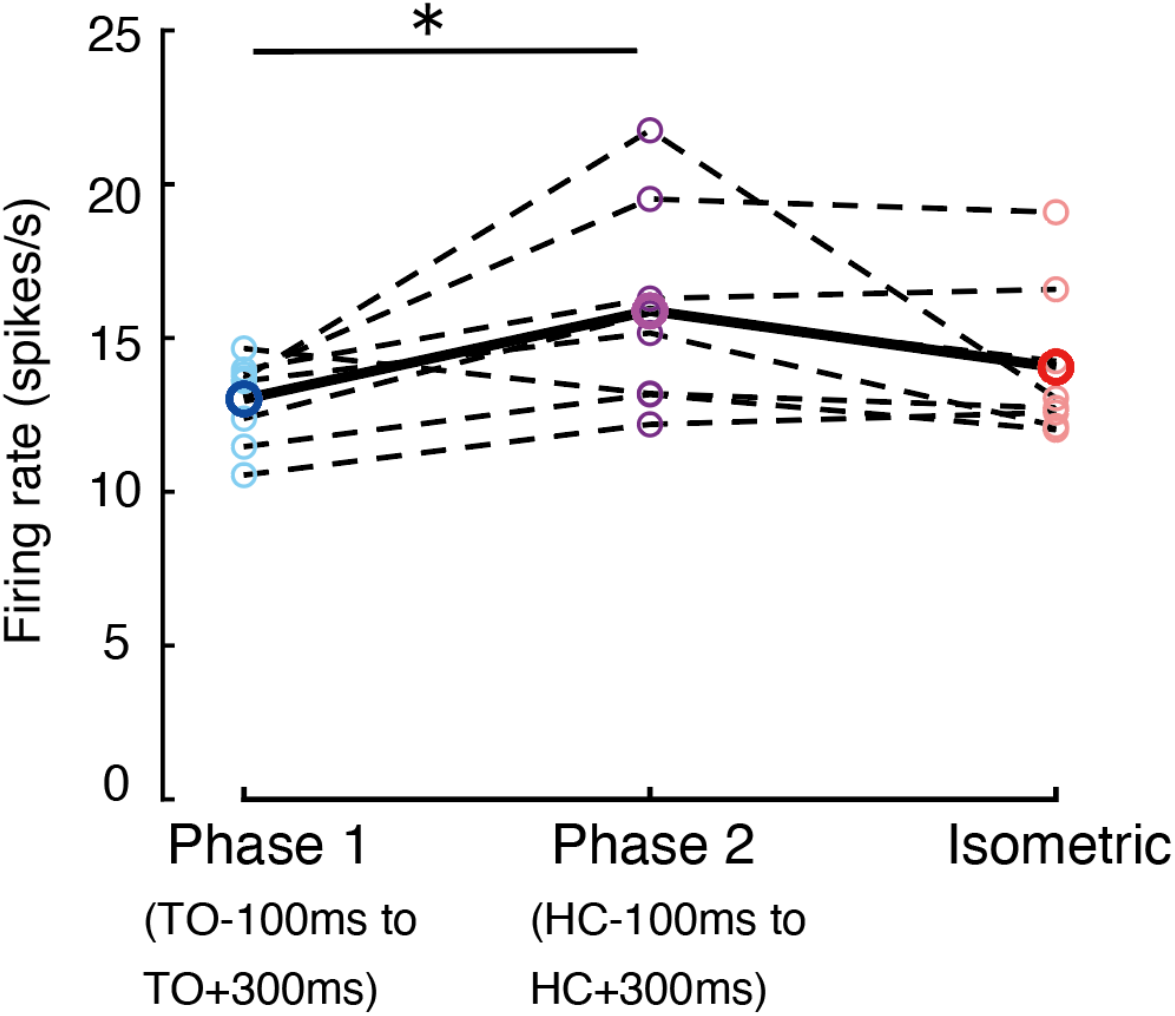
Firing rate of motor neuron discharges. Mean (solid line with thick-lined circles) and individual (dashed line with thin-lined circles) values are shown. * p <0.05 (permutation test with FDR correction).

## Discussion

This study is the first to show MN firing activity during human walking using HDsEMG decomposition. Ten to twenty MUs were identified in most participants (Fig. 5). We found doublet firing, activation phase specificity, and MU recruitment modulation to be features of MU firing behavior during walking (Table 1; Figs. 6–7). We also observed high MN synchronization across a wide range of frequencies (Figs. 8–9). These results greatly advance our understanding of neuromuscular control during walking.

### Doublet firings during walking

Some MNs exhibited doublets during walking (Table 1) and were associated with peak TA activity (Fig. 6). Doublets significantly enhance muscle force output (Mrówczyński et al., 2015). The enhancement was two to three times greater than that by a single firing (Kamavuako and Farina, 2010). Doublets are frequently observed when muscle force increases rapidly rather than during tonic contractions (Mrówczyński et al., 2015).

In the present study, doublets frequently occurred during three peak timings of TA activity: pre-swing, mid-swing, and loading response phases (Fig. 6). Initial stance doublets probably generate ankle dorsiflexion force to resist ankle joint rotation in the plantarflexion direction after heel contact (Fig. 6). The doublets in the two timings of the swing phase are considered to contribute to: (1) preventing excessive ankle joint extension by eccentric contraction in the pre-swing phase and (2) providing toe clearance by concentric contraction mid-swing.

Based on existing knowledge of their possible functions, doublets may be a central nervous system strategy to efficiently control muscle force during gait. Doublets are related to the strength of synaptic inputs to MNs (Gorassini et al., 2000; Krutki et al., 2010). Therefore, the rapid increase in synaptic inputs by a descending drive or afferent feedback evokes doublets during walking. Supporting this hypothesis, significant functional connectivity between the motor cortex and TA has been reported at pre-swing (Roeder et al., 2018; Yokoyama et al., 2020), mid-swing (Petersen et al., 2012), and initial stance (Roeder et al., 2018; Yokoyama et al., 2020). Additionally, loading afferent feedback during the initial stance phase plays a crucial role in generating TA activity (Dietz, 2002a).

### MU recruitment during walking

We found differences in the MU recruitment threshold between isometric contraction and walking tasks (Fig. 7). In general, MU recruitment during voluntary contractions follows an order consistent with the size principle (Henneman et al., 1965). However, some exceptions have been reported in humans during voluntary contractions. For example, MU recruitment order can differ depending on motor task (Thomas et al., 1986; Hudson et al., 2017). Hudson et al. (2017) reported different MU recruitment patterns of the parasternal intercostal muscle during voluntary trunk rotation and involuntary breathing. Because walking control consists of voluntary and involuntary components (Zijlstra et al., 1995), differences in involuntariness between walking and isometric contractions, which are more voluntary, may be related to the recruitment threshold modulations in the two tasks. In addition, afferent feedback to the MN pool can affect the MU recruitment threshold (Garnett and Stephens, 1981). Compared with isometric contractions, walking is associated with different sensory inputs, such as proprioceptive afferents from joint movements and load afferent inputs from the sole. Differences in sensory information may also affect MU recruitment threshold modulation.

We also observed activation phase-specific MU recruitment during walking in some MUs (Table 1). The number of participants with phase 1-specific MUs was limited (38.5%, Table 1). However, considering the few phase 1-specific MUs within all identified MUs (9.20%, Table 1), it is possible that phase 1-specific MUs existed in most participants but were not identified by HDsEMG decomposition because it extracts very few parts of the entire muscle’s MUs (Farina and Holobar, 2016). Such phase-specific recruitment of MUs was also demonstrated, albeit in a small number of MU populations, in a study using intramuscular EMG during walking with limited ankle motion (De Serres et al., 1995). TA activation during the swing and stance phases is functionally distinct (Perry, 1992). Given the task specificity of MU recruitment in TA (Hudson et al., 2019), functional differences may be involved in the activation phase-specific MU recruitment during walking. Regarding sensory information during walking, substantial load afferent inputs arise during the stance phase (Nilsson and Thorstensson, 1989) and proprioceptive information differs between the stance and swing phases (Dietz, 2002b). Therefore, sensory input differences between swing and stance phases may also be related to activation phase-specific MU recruitment.

### Highly synchronized firings during walking

We found highly synchronized firing during walking by time-and frequency-domain analyses (Figs. 8–9). Significant coherence was observed in broader frequencies below 50 Hz, especially below 5 Hz. MN synchronization strength is likely related to the amount of common synaptic inputs to the MN pool (Farina et al., 2016). Low-frequency MN synchronization components (<5 Hz) play a role in muscle force control (Farina and Negro, 2015). In the present study, the low-frequency component was dominant in MN synchronization during walking (Fig. 9A). MN synchronization in the alpha band (7.5–10 Hz) has been suggested to be related to afferent feedback (Erimaki and Christakos, 1999) and movement speed (Wessberg and Kakuda, 1999). Higher-frequency components in MU synchronization (12.5–40 Hz) are generally considered to be of cortical origin (Farina et al., 2014). During walking, a peak of coherence was observed in the beta band (15–40 Hz), in addition to high coherence in the lower frequency component (<5 Hz) (Fig. 9A). In neonatal leg movements, spinal CPGs cause high MN synchronization in the low-frequency band (Del Vecchio et al., 2020). Considering the shared primitive motor circuits in leg movements between human infants and adults (Dominici et al., 2011), the spinal CPG would contribute to the high MN synchronization in the low-frequency bandwidth in adult walking. In addition to the spinal CPGs, the cortex is probably also involved in MN synchronization based on beta band peak coherence (Fig. 9A). The cortical and spinal contributions to the control of MNs correspond to current knowledge on the neural control of human locomotion (Nielsen, 2016).

The coherence differed between phase 1 and 2 of TA activity during walking (Fig. 9B). High-beta (22.5–40 Hz) cortical activity is involved in muscular control when motor preparation is required (Salenius et al., 1996). Thus, descending inputs from the cortex may contribute to high MN synchronization in the high-beta band in phase 2 (Fig. 9B), providing predictive control of the TA to improve the loading response after heel contact (Ogawa et al., 2014). The lower frequency band MN synchronization (<5 Hz) was also higher during phase 2 than phase 1 (Fig. 9B). Because cutaneous foot afferents affect MN activity in the TA (Nielsen et al., 1997), we speculate that strong afferent inputs from the sole at the initial stance may cause high MN synchronization, which is supported by a study suggesting that sensory afferents contribute to MN synchronization during infant leg movements (Del Vecchio et al., 2020). Additionally, the higher firing rate of MUs in phase 2 than phase 1 (Fig. 10) probably contributed higher synchronization (Fig. 9) because it is positively affected by the average firing rate (De La Rocha et al., 2007; Vecchio et al., 2019). However, because the average firing rates were generally similar between phases 1 and 2 (13.0 and 15.9 spikes/s, respectively) compared to the firing rate ranges tested previously (20-80 spikes/s) (Vecchio et al., 2019), the effect of firing rate on MU synchronization may not have been dominant.

We also found that MN synchronization was generally higher during walking than during isometric contraction (Figs. 8–9). This difference may be derived from a suggested function that MU synchronization contributes to force development during rapid contractions (Semmler, 2002). High MN synchronization at low frequency (<5 Hz) during walking compared to isometric contraction (Fig. 9B) may be due to the involvement of primitive locomotor circuits (Grillner, 2011), consistent with the higher MN synchronization in infants’ leg movements than in isometric contractions in adults (Del Vecchio et al., 2020). Higher alpha-band MN synchronization in phase 2 than in isometric contraction (Fig. 9B) was probably related to a positive correlation between alpha-band MN synchronization and movement speed (Wessberg and Kakuda, 1999). We also found higher MN synchronization in the beta band during walking compared to isometric contractions (Fig. 9B). Beta-band corticomuscular coherence is positively correlated with force variability (Ushiyama et al., 2017). Since the isometric contraction task required stable force output, beta-band MU synchronization during isometric contractions was possibly low to prevent high variability in muscle force.

### Conclusion

This is the first study exploring MU firing behavior during walking by HDsEMG decomposition of non-invasive, wireless EMG recordings to minimize movement artifacts and revealed MN firing behavior regarding doublets, recruitment patterns, and synchronization. The results suggest that the central nervous system, including the spinal CPG and cerebral cortex, flexibly controls MN firing to generate the muscle force required during walking. These novel findings greatly expand our understanding of the neural control of walking and will be fundamental for future studies on individuals with gait disorders.

## Acknowledgments

The work of Hikaru Yokoyama was supported by the Grant-in-Aid for Scientific Research (B) through Japan Society for the Promotion of Science (JSPS) under Grant 21H03340.

We thank Dr. Moeka Yokoyama for her helpful comments and discussion.

